# Insights into the molecular mechanism of translation inhibition by the ribosome-targeting antibiotic thermorubin

**DOI:** 10.1101/2022.09.15.508020

**Authors:** Madhura N. Paranjpe, Valeria I. Marina, Aleksandr A. Grachev, Tinashe P. Maviza, Olga A. Tolicheva, Alena Paleskava, Ilya A. Osterman, Petr V. Sergiev, Andrey L. Konevega, Yury S. Polikanov, Matthieu G. Gagnon

## Abstract

Thermorubin (THR) is an aromatic anthracenopyranone antibiotic active against both Gram-positive and Gram-negative bacteria. It is known to bind to the 70S ribosome at the intersubunit bridge B2a and was thought to inhibit factor-dependent initiation of translation and obstruct the accommodation of tRNAs into the A site. Here, we show that thermorubin causes ribosomes to stall *in vivo* and *in vitro* at internal and termination codons, thereby allowing the ribosome to initiate protein synthesis and translate at least a few codons before stalling. Our biochemical data show that THR affects multiple steps of translation elongation with a significant impact on the binding stability of the tRNA in the A site, explaining premature cessation of translation. Our high-resolution crystal and cryo-EM structures of the 70S-THR complex show that THR can co-exist with P- and A-site tRNAs, explaining how ribosomes can elongate in the presence of the drug. Remarkable is the ability of THR to arrest ribosomes at the stop codons. Our data suggest that by causing structural re-arrangements in the decoding center, THR interferes with the accommodation of tRNAs or release factors into the ribosomal A site.

**HIGHLIGHTS:**

- Thermorubin is a potent inhibitor of protein synthesis both *in vivo* and *in vitro*;
- Thermorubin does not prevent the binding of P- and A-site tRNAs;
- Thermorubin affects multiple steps of translation elongation with a major impact on binding stability of the A-site tRNA;
- Thermorubin can act as an inhibitor of translation termination on some ORFs.

## INTRODUCTION

The ever-growing antibiotic resistance among bacterial pathogens represents a major threat to public health care (1), calling for the continued search for new and improvements of existing drugs. Understanding the detailed molecular mechanisms by which various antimicrobials impede certain cellular functions is critical for their potential improvement to better suit clinical applications and circumvent development of resistance. About half of the antibiotics currently used in the clinic slow down or kill pathogenic bacteria and thereby cure infections by selectively inhibiting their ribosomes – the central components of the protein synthesis apparatus. Being the most conserved and sophisticated molecular machine of the cell, the ribosome is composed of two unequal separable subunits, small and large (30S and 50S in bacteria), which join together to form the 70S ribosome (2). Genetic information for protein synthesis is carried to the ribosome by messenger RNA (mRNA), which is read by nucleotide triplets (codons) with the help of auxiliary transfer RNA (tRNA) molecules that have two functional ends, one carrying the amino acid and the other containing the anticodon that recognizes the mRNA codons. tRNAs are indispensable adaptors that bridge the two main functional centers of the ribosome: the decoding center (DC) in the small subunit and the catalytic peptidyl transferase center (PTC) at the heart of the large subunit. While the PTC chemically links amino acids delivered by the tRNAs into polypeptides, the DC monitors the accuracy of base pairing between the codon of the mRNA and the anticodon of the tRNA, ensuring that the correct tRNAs are selected. The majority of ribosome-targeting antibiotics interfere with protein synthesis by binding at the ribosome functional centers (such as DC or PTC) and either freeze a particular conformation of the ribosome or hinder the binding of its ligands (3). In particular, DC is targeted by many chemically unrelated antibiotics, including aminoglycosides, tetracyclines, tuberactinomycins, odilorhabdins, and negamycin (3,4).

In 1964, a secondary metabolite from *Thermoactinomyces antibioticus*, called thermorubin (THR), was shown to exhibit a broad range of activity against both Gram-positive and Gram-negative bacteria (5,6). THR was further shown to inhibit protein synthesis *in vitro* (7), suggesting that the ribosome might be its primary intracellular target. Only much later, in 2012, the first structure of a vacant 70S ribosome from the Gram-negative bacterium *Thermus thermophilus* in complex with THR (8) revealed that, despite being structurally similar to tetracyclines, THR binds to a distinct site within DC largely overlapping with those of aminoglycosides and tuberactinomycins (9,10). The binding site of THR is located at the interface between the 30S and 50S ribosomal subunits within bridge B2a (8). The tetracyclic moiety of THR stacks between residue A1913 of Helix 69 (H69) of the 23S rRNA and the C1409-G1491 basepair at the top of helix 44 (h44) of the 16S rRNA, whereas the orthohydroxyphenyl moiety of THR stacks upon the nucleobase of U1915 in H69 (8). Importantly, THR does not affect translation in eukaryotic cells (5,6), such as yeast and fungi, which can be rationalized by the absence of the equivalent base pair in the 18S rRNA and the concomitant inability to stack with the tetracyclic moiety of the drug.

The precise mechanism of THR action during protein synthesis remains unclear. Early biochemical studies suggested that THR inhibits binding of initiator tRNA to the P site in a factor-dependent manner (7,11). Initiation factors 1 (IF1) and 2 (IF2) interact with bridge B2a, suggesting that, in principle, ribosome-bound THR could interfere with the conformational changes of this region required for efficient delivery of initiator fMet-tRNA to the P site during translation initiation (8). However, from the previous structure, it became evident that the THR binding site in the vacant ribosome does not directly overlap with any of the tRNA binding sites (8). Nevertheless, the interaction of THR with bridge B2a induces nucleotide C1914 to flip out of H69, taking a conformation that is incompatible with the binding of an aminoacyl-tRNAs into the A site (8). Thus, contrary to the concept of being an initiation inhibitor (7), Bulkley *et al*. argued that THR is likely to interfere with the accommodation of an aminoacyl-tRNAs to the A site, including the very first round of translation elongation, which could be considered as inhibition of translation initiation (8). However, the progression and stalling of translating ribosomes along mRNA in the presence of THR have never been assessed. Moreover, given that many of the 16S and 23S rRNA nucleotides, especially those constituting DC and the B2a bridge, change conformation upon tRNA binding, it is conceivable that the mode of THR binding to the ribosome could be different in the presence of native tRNA substrates. Finally, it is difficult to reconcile how a single nucleotide (C1914) located in a relatively unconfined pocket of the ribosome having multiple opportunities for deflection to avoid a clash with the A-site tRNA, can prevent binding of a much larger tRNA molecule to the ribosomal A site.

In this work, we set to close these gaps and uncover the molecular mechanism underlying the mode of THR inhibition of translation. Using a combination of *in vivo* and *in vitro* techniques, we show that THR targets bacterial ribosomes, and instead of arresting the ribosome at the start codon, it does so at various codons along mRNA templates. Remarkably, on some mRNAs, THR stalls the ribosome at the stop codon. By solving the 2.7Å resolution X-ray crystal structure of THR in complex with the *T. thermophilus* 70S ribosome carrying initiator formyl-methionyl-tRNA_i_^fMet^, we demonstrate that neither the binding site of the ribosome-bound drug nor the location of the P-site tRNA is affected by the presence of each other. We further determined the 2.7Å resolution cryo-EM structure of the *Escherichia coli* 70S ribosome in complex with the drug and carrying both A- and P-site tRNAs, in line with our biochemical data suggesting that THR can co-exist on the ribosome with the A-site tRNA. Furthermore, we show that multiple partial reactions of the translation elongation cycle are affected by THR, with the most pronounced effect observed on the binding stability of A-site tRNA, explaining the internal ribosome stalling events. Our structural data also suggest that THR can arrest ribosomes at termination codons likely by preventing accommodation of release factor. Altogether, our results provide a molecular understanding of the mode of ribosome inhibition by THR.

## MATERIALS AND METHODS

### Reagents

All reagents and chemicals were obtained from MilliporeSigma (USA). Thermorubin was isolated from *Thermoactinomyces antibioticus* as reported previously (5) and provided by Dr. Francis Johnson.

### In vivo detection of translation inhibitors using the pDualrep2 reporter system

For the *in vivo* bioactivity test (**Figure 1A**), we used the reporter strain BW25113ΔtolC-pDualrep2 as described previously (12,13). Briefly, 1.5 µL of the solutions of THR (10 mg/ml), erythromycin (ERY, 5 mg/ml), and levofloxacin (LEV, 25 µg/ml) in DMSO were applied onto the agar plate that already contained a lawn of the reporter strain. After overnight incubation at 37°C, the plate was scanned by ChemiDoc MP (Bio-Rad, USA) using “Cy3-blot” mode for RFP fluorescence and “Cy5-blot” mode for Katushka2S fluorescence.

**Figure 1.**
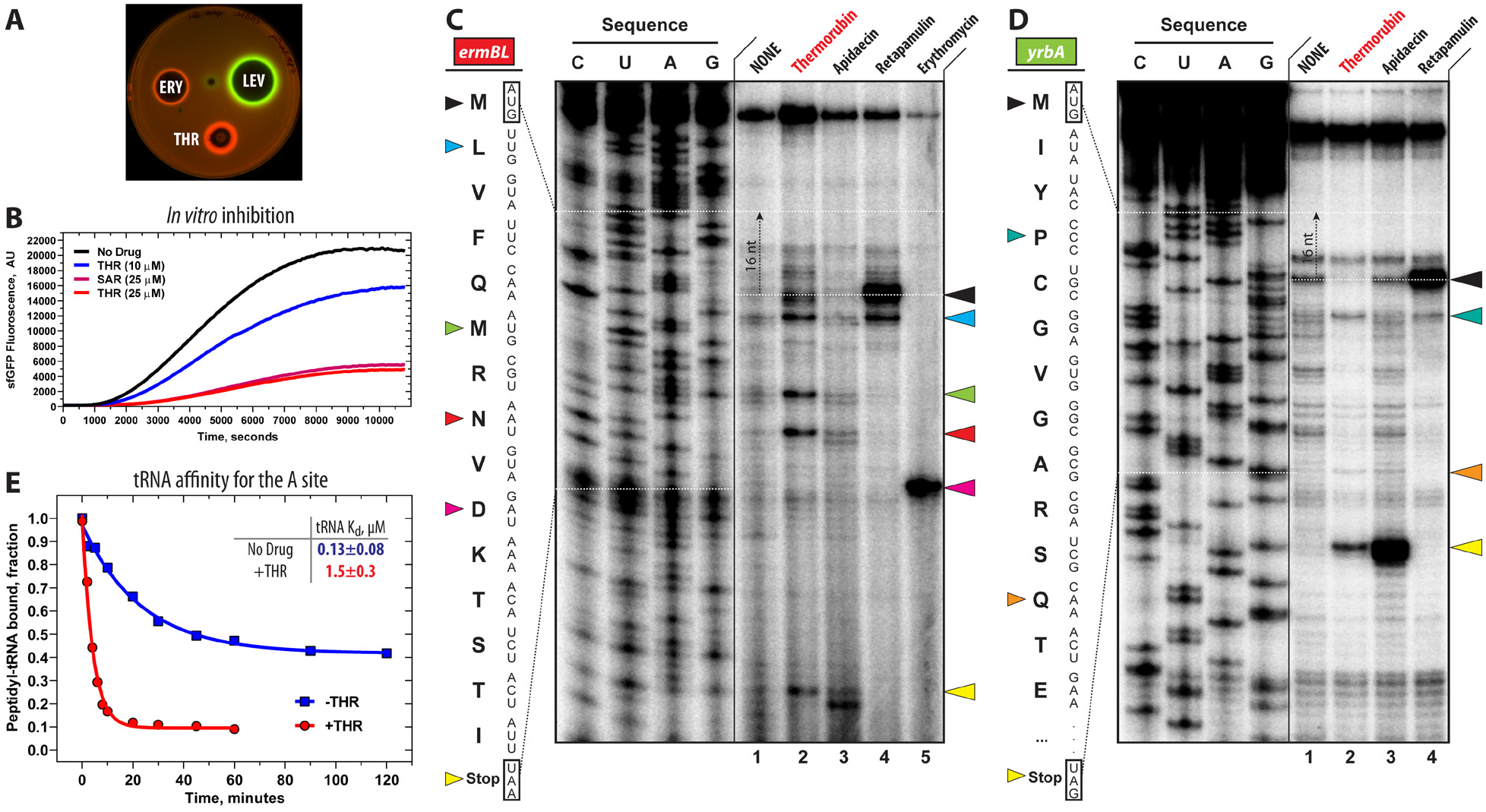
Thermorubin inhibits protein synthesis both *in vivo* and *in vitro*. (**A**) Induction of a two-color dual reporter system sensitive to inhibitors of the ribosome progression or inhibitors of DNA replication. Spots of erythromycin (ERY), levofloxacin (LEV), and thermorubin (THR) were placed on the surface of an agar plate containing *E. coli* ΔtolC cells transformed with the pDualrep2 plasmid (13). Induction of expression of Katushka2S (red) is triggered by translation inhibitors, while RFP (green) is induced upon DNA damage. (**B**) Time-courses of inhibition of sfGFP synthesis by THR (blue and red curves) and sarecycline (SAR, magenta curve) in the *in vitro* cell-free translation system. AU, arbitrary units. (**C, D**) Ribosome stalling by THR and other protein synthesis inhibitors on *ermBL* (C) and *yrbA* (D) mRNA templates as revealed by reverse transcription inhibition (toe-printing) in a recombinant cell-free translation system. Nucleotide sequences of wild-type *ermBL* and *yrbA* genes, and the corresponding amino acid sequence, are shown on the left in each panel. Sequencing lanes (C, U, A, G) are on the left of each gel. Due to the large size of the ribosome, the reverse transcriptase used in the toe-printing assay stops 16 nucleotides downstream of the codon located in the P site. Black arrowhead marks the translation start site (retapamulin control, lane 4). Blue, green, red, teal, and orange arrowheads point to the THR-induced arrest sites within the coding sequences of *ermBL* or *yrbA* mRNAs. Magenta arrowhead points to the erythromycin-specific stalling site on *ermBL* only. Yellow arrowheads point to the translation stop site (apidaecin control, lane 3). Note that, similar to apidaecin, THR induces ribosome stalling at the stop codons of *ermBL* and *yrbA* mRNAs. Experiments were repeated twice independently with similar results. (**E**) Time courses of dissociation of peptidyl-tRNA from the A site of the 70S pre-translocation ribosome complexes in the absence (blue) or presence (red) of THR.

### In vitro translation inhibition assay

To test the ability of THR to inhibit protein synthesis *in vitro* (**Figure 1B**), we used the PURExpress translation system (New England Biolabs) reconstituted from purified components. Translation of superfolder green fluorescent protein (sfGFP) was carried out according to the manufacturer’s protocol. The assembled reactions (5 µL) were supplemented with 5 ng of a plasmid encoding sfGFPRNA. Varying concentrations of THR (0 μM, 10 μM, or 25 μM) or sarecycline (SAR, 25 μM) were added to the reaction mixtures, which were then placed in a 384-well black-wall plate, and the progression of each reaction was monitored over 3 hours at 37°C by a TECAN microplate scanner with the excitation and emission wavelengths set at 488 nm and 520 nm, respectively.

### Toe-printing analysis

The synthetic DNA templates encoding the amino acid sequences for Rst1, Rst2, ErmBL, and YrbA open reading frames (ORFs) were generated by polymerase chain reaction (PCR) and AccuPrime Taq DNA Polymerase (Thermo Fisher Scientific, USA). The sequences of the primers used for PCR are shown in **Table S1**. The toe-printing analysis of drug-dependent ribosome stalling analysis (**Figure 1C, D; Figure S1**) was carried out using Rst1, Rst2, ErmBL, and YrbA mRNA templates as previously described (14,15) with minor modifications. Toe-printing reactions were carried out in 5-µl aliquots containing a PURExpress transcription-translation coupled system (New England Biolabs, USA) to which the test template was added (16). The reactions were incubated at 37°C for 20-25 minutes. Reverse transcription on the templates was carried out using radioactively labeled primer NV1 (**Table S1**). Primer extension products were resolved on 6% sequencing gels as described previously (17). The final concentrations of drugs were: 50 µM THR, 50 µM apidaecin (API), 50 µM retapamulin (RET), and 50 µM erythromycin (ERY).

### X-ray crystallographic structure determination

Wild-type 70S ribosomes from *T. thermophilus* (strain HB8) were prepared as described previously (18-21). Synthetic mRNA with the sequence 5’-GGC-AAG-GAG-GUA-AAA-**AUG**-**UAA**-3’ (24MStop) containing Shine-Dalgarno sequence followed by the P-site methionine and the A-site stop codons was obtained from Integrated DNA Technologies (Coralville, IA, USA). Wild-type deacylated initiator tRNA_i_^fMet^ was overexpressed and purified from *E. coli* as described previously (22-25). The complex of the wild-type *T. thermophilus* 70S ribosome with mRNA, vacant A site, and P-site tRNA_i_^fMet^ was formed as described previously (20,21,23) in the presence of 500 µM THR.

Collection and processing of the X-ray diffraction data, model building, and structure refinement were performed as described in our previous reports (20,21,23,24,26,27). Diffraction data were collected at beamlines 24ID-C and 24ID-E at the Advanced Photon Source (Argonne National Laboratory). A complete dataset for each complex was collected using 0.979 Å irradiation at 100 K from multiple regions of the same crystal, using 0.3-degree oscillations. Raw data were integrated and scaled using the XDS software package (Feb 5, 2021) (28). Molecular replacement was performed using PHASER from the CCP4 program suite (version 7.0) (29). The search model was generated from the previously published structures of the *T. thermophilus* 70S ribosome with bound mRNA and aminoacylated tRNAs (PDB entry 6XHW (23)). Initial molecular replacement solutions were refined by rigid-body refinement with the ribosome split into multiple domains, followed by positional and individual B-factor refinement using the PHENIX software (version 1.17) (30). Non-crystallographic symmetry restraints were applied to four parts of the 30S ribosomal subunit (head, body, spur, and helix 44) and four parts of the 50S subunit (body, L1-stalk, L10-stalk, and C-terminus of the L9 protein). Structural models were built in Coot (version 0.8.2) (31). The statistics of data collection and refinement are compiled in **Table S2**.

### Sample preparation for cryo-EM, data acquisition, and image processing

The *E. coli* 70S ribosomes, isolated from strain MRE600, were prepared as previously described with some modifications (32). Briefly, the cells were washed in buffer containing 20 mM Tris-HCl pH 7.4, 10.5 mM MgCl_2_, 100 mM NH_4_Cl, 0.5 mM EDTA, 6 mM β-mercaptoethanol, and DNase I (16 U/g cells) and then lysed using a microfluidizer LM20-30 (Microfluidics, Westwood, MA). The lysate was cleared by centrifugation at 16,000 rpm (30,600 × *g*) for 1 hour at 4°C and filter-sterilized through a 0.45 μm filter. To remove fine particles and residual debris from the supernatant, the lysate was spun at 30,000 rpm (104,000 × *g*) in a Type 45Ti rotor (Beckman) for 30 min at 4°C. To isolate ribosomes, the lysate was layered on a 1.1 M sucrose cushion buffer (20 mM Tris-HCl pH 7.4, 500 mM NH_4_Cl, 10.5 mM MgCl_2_, 0.5 mM EDTA, 6 mM β-mercaptoethanol) and spun at 43,000 rpm (214,000 × *g*) in a Type 45Ti rotor (Beckman) for 19 hours at 4°C. Ribosome pellets were resuspended in 20 mM Tris-HCl pH 7.4, 10.5 mM MgCl_2_, 100 mM NH_4_Cl, 0.5 mM EDTA and 6 mM β-mercaptoethanol. Ribosomes were then purified through 10-40% sucrose density gradients in a SW32 rotor (Beckman) at 23,000 rpm (90,000 × *g*) at 4°C for 13 hours. The fractions containing 70S ribosomes were collected, diluted to adjust Mg^2+^ concentration to 10 mM, and concentrated by centrifugation at 43,000 rpm (214,000 × *g*) at 4°C. Pure 70S ribosomes were resuspended and brought to the final buffer containing 10 mM Tris-HCl pH 7.4, 10 mM MgCl_2_, 60 mM NH_4_Cl, and 6 mM β-mercaptoethanol, flash-frozen in liquid nitrogen and stored at –80°C.

To determine the cryo-EM structure of the *E. coli* 70S-THR complex with tRNAs, we incubated 2 µM *E. coli* 70S ribosomes, 8 µM 24-MF mRNA, 8 µM fMet-tRNA_i_^fMet^ in 1x ribosome buffer (5 mM Tris-HCl pH 7.4, 60 mM NH_4_Cl, 10 mM MgCl_2_, 6 mM β-mercaptoethanol) at 37 °C for 10 min. Then, 200 µM THR was added and the complex was incubated at room temperature for 10 minutes. Finally, 15 µM Phe-tRNA^Phe^ was added and incubated for an additional 10 minutes at room temperature.

Quantifoil R2/1 gold 200 mesh grids (Electron Microscopy Sciences) were glow-discharged for 30 s in an (H_2_O_2_)-atmosphere using the Solarus 950 plasma cleaner (Gatan). The mixture (4 μl), containing 2 μM *E. coli* 70S-THR carrying A- and P-site tRNAs, was applied onto grids, blotted in 85% humidity at 22 °C for 18 s, and plunged-frozen in liquid nitrogen-cooled ethane using a Leica EM GP2 cryo-plunger. Grids were transferred into a Titan Krios G3i electron microscope (ThermoFisher Scientific) operating at 300 keV and equipped with a Falcon3 direct electron detector camera (ThermoFisher Scientific). The image stacks (movies) were acquired with a pixel size of 0.86 Å/pixel. Data collection was done in the EPU software (ThermoFisher Scientific) setup to record movies with 39 fractions with a total accumulated dose of 40 e-/Å^2^/movie. A total of 10,032 image stacks were collected with a defocus ranging between –1 to –2.3 µm. The statistics of data acquisition are summarized in **Table S3**.

Data processing was done in cryoSPARC 3.3.2 (33). The image stacks were imported into cryoSPARC and gain-corrected. Image frames (fractions) were motion-corrected with dose-weighting using the patch motion correction, and patch contrast transfer function (CTF) estimation was performed on the motion-corrected micrographs. Based on relative ice thickness, CTF fit, length, and curvature of motion trajectories, 9,852 micrographs were selected for further processing (**Figure S4**).

982,190 particles were picked using the circular “blob” picker in cryoSPARC and were filtered based on defocus adjusted power and pick scores to 972,828 particles. Then, particles were subjected to two rounds of reference-free two-dimensional (2D) classification. After discarding bad particles, 613,586 particles were selected from 2D classification and used to generate the *ab-initio* volume. Using ‘heterogeneous refinement’ in cryoSPARC with five groups, particles were further classified into two class averages. Both class averages represent the 70S ribosome with density for bound tRNAs. Particles were therefore combined (565,280) and classified based on focused 3D variability analysis (3DVA) with a mask around the A-site tRNA which allowed to remove 55,534 “bad” particles with broken density for tRNAs. This approach yielded two main class averages containing 310,702 ribosome particles with A- and P-site tRNAs, and 199,044 ribosome particles containing P-site tRNA and no A-site tRNA. Particles from each class were re-extracted to full-size (512×512-pixel box), followed by non-uniform and CTF refinement in cryoSPARC. The Fourier Shell Correlation (FSC) curves were calculated using the cryo-EM validation tool in Phenix 1.19.2 for even and odd particle half-sets masked with a ‘soft mask’ excluding solvent (34). The nominal resolution of the reconstructions using the FSC-cutoff criterion of 0.143 is 2.7Å for the 70S-THR ribosome complexed with A-site Phe-tRNA^Phe^ and deacylated P-site tRNA_i_^fMet^, and 2.8Å for the 70S-THR ribosome complexed with P-site tRNA_i_^fMet^ (**Figure S5**).

### Model building and refinement of the E. coli 70S ribosome

As a starting model, the 30S and 50S subunits were taken from the structure of the *E. coli* 70S ribosome (PDB entry 6GXP (35)) and rigid-body docked into the 2.7Å-resolution cryo-EM map of the 70S-THR with A- and P-site tRNAs using UCSF Chimera 1.14 (36). The fMet-tRNA_i_^fMet^ and Phe-tRNA^Phe^ were taken from PDB entry 6XHW (23), rigid-body fit into the EM density, and adjusted in Coot (37). Based on the high-resolution structure of the *E. coli* 70S ribosome (PDB entry 7K00 (38)), modified nucleotides were included in the model. The complete model of the *E. coli* 70S ribosome with ordered solvent and bound THR, A-site Phe-tRNA^Phe^ and P-site tRNA_i_^fMet^ was real-space refined into the EM map for five cycles using Phenix 1.19.2 (39) including global energy minimization and group ADP refinement strategies along with base pair restraints for rRNA and tRNAs, together with Ramachandran and secondary structure restraints. The resulting model of the *E. coli* 70S-THR with Phe-tRNA^Phe^ in the A site and deacylated tRNA_i_^fMet^ in the P site was validated using the comprehensive validation tool for cryo-EM in Phenix 1.19.2 (34). The cryo-EM data collection, refinement, and validation statistics are compiled in **Table S3**.

### Figure generation

All figures showing atomic models were rendered using the PyMol software (www.pymol.org), UCSF Chimera 1.14 (36), or Chimera X 1.2 (40) and assembled with Adobe Illustrator (Adobe Inc.).

### Sample preparation for rapid kinetics measurements

*E. coli* 70S ribosomes, initiation factors (IF1, IF2, and IF3), elongation factors (EF-Tu and EF-G), aminoacylated initiator fMet-tRNA_i_^fMet^, aminoacylated [^14^C]Phe-tRNA^Phe^, tRNA^Phe^(Prf16/17), and [^14^C]Val-tRNA^Val^ were prepared as described previously (41-44). MF-mRNA containing AUG-UUU nucleotide sequence encoding for Met-Phe dipeptide and MVF-mRNA containing AUG-GUU-UUU nucleotide sequence encoding for Met-Val-Phe tripeptide were obtained by T7 transcription. The full sequences for the DNA templates used for T7 transcription of MF- and MVF-mRNAs are:

- MF-mRNA: CGAATTTA*ATACGACTCACTATAG*GGAATTCAAAAATTTAAAAGTTAACAGGTATA CATACT**ATGTTT**ACGATTACTACGATCTTCTTCACTTAATGCGTCTGCAGGCATG CAAGC
- MVF-mRNA: CGAATTTA*ATACGACTCACTATAG*GGAATTCAAAAATTTAAAAGTTAACAGGTATA CATACT**ATGGTTTTT**ATTACTACGATCTTCTTCACTTAATGCGTCTGCAGGCATG CAAC

The sequences of T7 promoters are italicized; sequences coding for Met-Phe or Met-Val-Phe are bold. All reactions were performed in TAKM7 buffer (50 mM Tris-HCl pH 7.5, 70 mM NH_4_Cl, 30 mM KCl, and 7 mM MgCl_2_) unless otherwise stated.

For a typical 1-ml reaction of initiation complex formation, 1 μM 70S ribosomes were incubated with 3 μM mRNA, 2 μM fMet-tRNA_i_^fMet^, 1.5 μM of IF1, IF2, and IF3 in TAKM7 buffer supplemented with 1 mM GTP and 2 mM DTT for 1 hour at 37°C. Ternary complexes of aminoacyl-tRNA*EF-Tu*GTP were prepared by preincubation of EF-Tu with 1 mM GTP, 3 mM phosphoenolpyruvate, 2 mM DTT, 1% pyruvate kinase in TAKM7 buffer for 15 minutes at 37°C, followed by addition of aminoacyl-tRNA and incubation for 1 minute. Pre-translocation ribosome complexes were formed by mixing 70S initiation and ternary complexes with subsequent incubation for 2 minutes at 37°C. For purification, the concentration of Mg^2+^ ions was increased to 21 mM, and the mixture of pre-translocation complexes was layered onto a 400-μl 1.1 M sucrose cushion (prepared in TAKM21 buffer), followed by centrifugation in an SW55 rotor (Beckman Coulter, USA) at 55,000 rpm for 3 hours at 4°C. The pellet was dissolved in TAKM21 buffer, flash frozen in liquid nitrogen, and stored at -80°C. fMet-tRNA_i_^fMet^, Phe-tRNA^Phe^, and Val-tRNA^Val^ were purified by HPLC and stored at -80°C. To prepare Phe-tRNA^Phe^(Prf16/17), 8 μM of deacylated tRNA^Phe^(Prf16/17) was incubated with 0.2 mM phenylalanine, 3 mM ATP, 6 μM β-mercaptoethanol, 40 nM phenylalanine-specific tRNA-synthetase, and 40 nM tRNA nucleotidyltransferase in TAKM7 buffer for 30 minutes at 37°C. Phe-tRNA^Phe^(Prf16/17) was purified on HiTrap Q column with 0.2 to 1 M NaCl gradient in 10 mM Tris-HCl pH 7.6, 10 mM MgCl_2_ and stored at -80°C. Where necessary, 33 μM THR (final concentrations after mixing are given throughout) was added to the 70S ribosomes, initiation, or pre-translocation complexes and incubated on ice for 2 minutes.

### Rapid kinetics measurements

To monitor the time course of aminoacyl-tRNA interaction with the A site of the ribosome, we rapidly mixed 0.1 µM Phe-tRNA^Phe^(Prf16/17)*EF-Tu*GTP ternary complex with 0.4 µM initiation ribosome complexes programmed with mRNA MF at 20°C. To study the pre-steady-state kinetics of translocation, we rapidly mixed 0.06 µM ribosome pre-translocation complexes containing deacylated tRNA_i_^fMet^ in the P site and fluorescently labeled fMet-Phe-tRNA^Phe^(Prf16/17) (45) in the A site with 2 µM EF-G in the presence of 1 mM GTP at 37°C. Fluorescence was recorded using SX-20 stopped-flow apparatus (Applied Photophysics). Proflavine fluorescence was excited at 460 nm and measured after passing through a KV495-nm cut-off filter. Light-scattering experiments were carried out at 430 nm, followed by detection at a 90° angle without the cut-off filter. Samples were rapidly mixed in equal volumes (60 µl). Time courses were obtained by averaging 5-to-7 individual transients (**Figure 5**). Data were evaluated by fitting with a two-exponential function with characteristic time constants (k_app1_, kapp2), amplitudes (A_1_, A_2_), and final signal amplitude (F_∞_) according to equation F = F_∞_ + A_1_*exp(-k_app1_*t) + A_2_*exp(-k_app2_*t), where F is the fluorescence at time t. All calculations were performed using GraphPad Prism 9.3.1 software (GraphPad Software, Inc).

### A-site peptidyl-tRNA dissociation constant measurements

Experiments and calculations were carried out according to the previously published methodology (46) in HAKM10 buffer (50 mM HEPES pH 7.5, 70 mM NH_4_Cl, 30 mM KCl, and 10 mM MgCl_2_) at 37°C. Briefly, to induce the dissociation of fMet-[^14^C]Val-tRNA^Val^ from the A site of purified pre-translocation complexes (0.25 μM), the Mg^2+^ concentration was adjusted from 21 to 10 mM in the sample, and the amount of peptidyl-tRNA bound to the A site was determined by nitrocellulose filtration at different incubation times. Dissociation elemental rate constant (*k*_*off*_) and equilibrium dissociation constant (K_d_) were calculated from time courses of dissociation by numerical integration (46). The following kinetic model was used: A ⇔ B + C, and B ⇒ D, where A denotes ribosomes with peptidyl-tRNA bound to the A site; B, unbound peptidyl-tRNA; C, ribosomes with unoccupied A site; D, hydrolyzed peptidyl-tRNA. The elemental association rate constant (*k*_*on*_) was calculated from the ratio K_d_ = *k*_*off*_/*k*_*on*_.

## RESULTS AND DISCUSSION

### Thermorubin targets translating ribosomes in vivo

The main goal of this study was to explore the inhibitory properties of anthracenopyranone antibiotic thermorubin (THR, **Figure 2A**) and uncover its mechanism of action. Although THR has long been known to inhibit protein synthesis in a crude cell lysate (7), it was also shown to exhibit potent activity against aldose reductase (47), suggesting that ribosome might not necessarily be the only cellular target for this drug. To check whether protein synthesis is the primary target of THR *in vivo*, we used an *E. coli*-based reporter system that is designed for rapid screening of inhibitors targeting either protein synthesis or DNA replication (13). In this assay, sub-inhibitory concentrations of antibacterial compounds that stall translation, such as the macrolide erythromycin (ERY), induce expression of the far-red fluorescent protein reporter Katushka2S (**Figure 1A**, ERY, red pseudocolor ring). Compounds that trigger the SOS response, such as inhibitors of DNA gyrase (e.g., levofloxacin (LEV)), induce expression of the Red Fluorescent Protein (RFP) reporter (**Figure 1A**, LEV, green pseudocolor ring). Both reporters are encoded by the same pDualerp2 plasmid in the *E. coli* BW25113 cells. The test antibiotic is applied as a drop or a disc onto a lawn of these cells, and its intracellular target could be determined based on the reporter being induced. When *E. coli* cells carrying the reporters were exposed to THR, the expression of the far-red fluorescent protein Katushka2S, but not of RFP, was observed, indicating that the drug specifically arrests ribosomes and targets protein synthesis *in vivo* (**Figure 1A**, THR, red pseudocolor ring).

**Figure 2.**
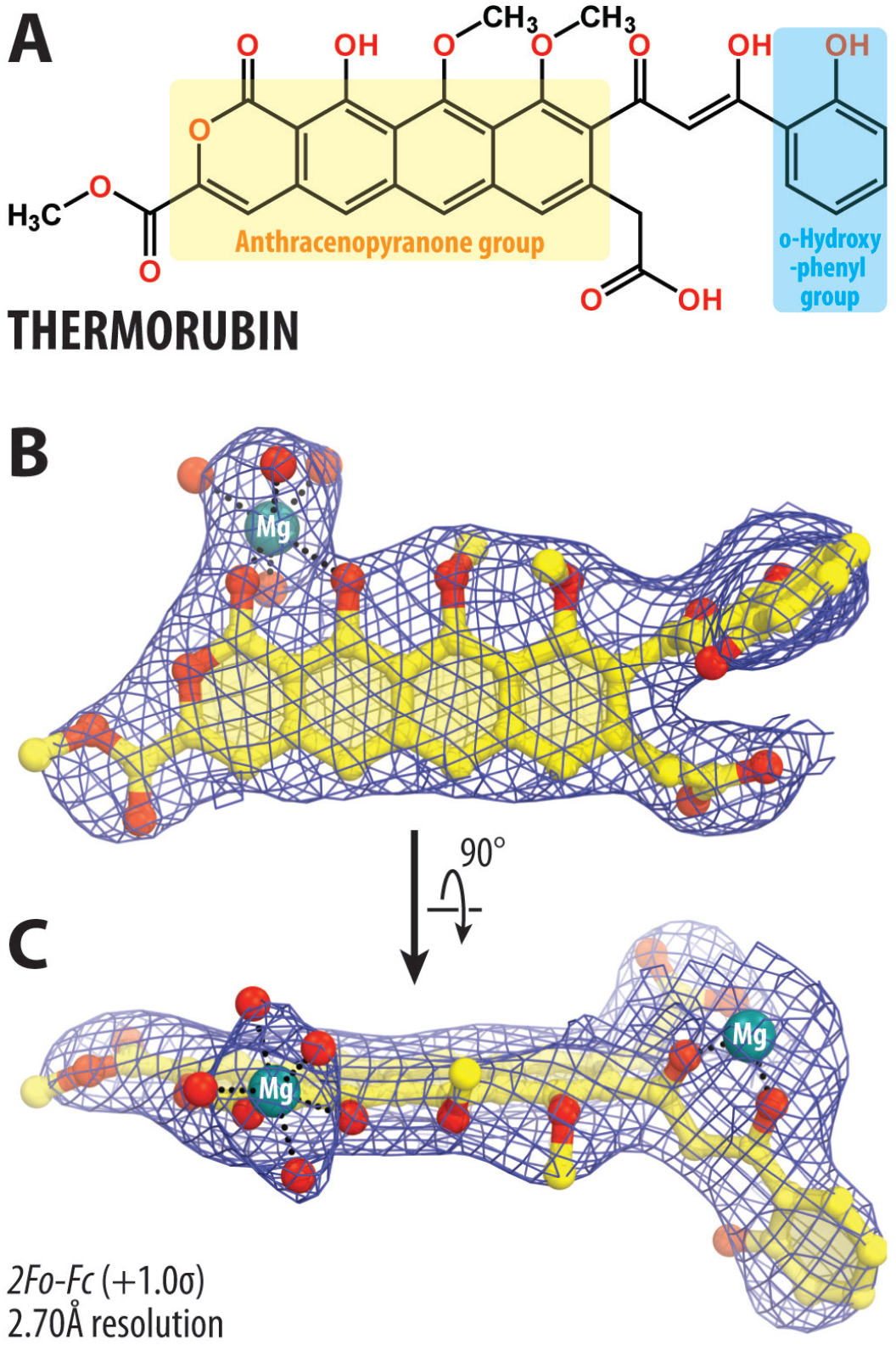
Chemical structure and X-ray electron density maps of ribosome-bound thermorubin. (**A**) Chemical structure of thermorubin (THR). (**B, C**) 2*Fo-Fc* Fourier electron density map of THR in complex with the *T. thermophilus* 70S ribosome (blue mesh) viewed from two different angles. The refined model of THR is displayed within its respective electron density map contoured at 1.0σ. Carbon atoms are colored yellow, oxygen atoms are red, and magnesium ions are teal. Note that the locations of all chemical moieties of THR can be unambiguously determined from the electron density map.

### Thermorubin is a potent inhibitor of protein synthesis in vitro

To further confirm that THR acts as a translation inhibitor, we tested its ability to interfere with *in vitro* protein synthesis. The addition of THR to the cell-free transcription-translation coupled system reconstituted from purified components resulted in a dose-dependent inhibition of the synthesis of superfolder green fluorescent protein (sfGFP) (**Figure 1B**). In these experiments, addition of 25 µM THR decreased sfGFP synthesis by ∼10 fold, comparable to the inhibitory properties of the tetracycline-class antibiotic sarecycline, which we used as a positive control. This data suggests that THR is a potent translation inhibitor and provides a promising foundation for future ribosome-targeting drug development.

It remains unclear, however, which step of translation – initiation, elongation, or termination – is inhibited by THR. Therefore, we used primer extension inhibition assay (toe-printing) to answer this question. This method is based on the reverse-transcriptase-dependent extension of a radioactively-labeled DNA primer annealed to the 3’-end of the translated mRNA template. Primer extension reaction terminates when the reverse transcriptase encounters a stalled ribosome. This technique allows us to unambiguously identify drug-induced ribosome stalling site(s) along the mRNA with single nucleotide precisions (15,48). A large set of ribosome-targeting antibiotics has been previously tested by toe-printing using *ermBL, yrbA, rst1*, or *rst2* mRNA templates (15) because their sequences contain codons for almost all proteinogenic amino acids. Therefore, we have chosen the same experimental system to directly compare our results for THR with those of other antibiotics with well-established modes of action.

The addition of THR to the cell-free translation system programmed with *ermBL, yrbA, rst1* or *rst2* mRNAs resulted in ribosome stalling at various codons within the corresponding open reading frame (ORF) but not at the start codons (**Figure 1C, D; Figure S1**). These data suggest that THR is not a *bona fide* initiation inhibitor, as speculated before (7). Interestingly, the ribosomes translate at least two codons before they stall in the presence of THR (**Figure 1C**, lane 2, blue arrowhead), excluding the possibility of THR acting during the first elongation cycle (at least on the mRNAs tested). Importantly, our experimental toe-printing data showing that THR allows for ribosome progression contradicts the previous structure-inspired idea that THR-dependent rearrangement of nucleotide C1914 prevents binding of the A-site tRNA and, thus, inhibits every elongation cycle (8). These data suggest that either THR can co-exist on the ribosome with an A-site tRNA, or it can bind and dissociate from the ribosome during the elongation and act in a context-specific fashion, i.e., preferably bind when certain codons are present in the A site. Perhaps, the most unexpected result of the toe-printing analysis was manifested on the *yrbA* mRNA template (and to a lesser extent on *ermBL* mRNA), where THR caused the most prominent ribosome stalling at the stop codon (**Figure 1C, D**, lane 2, yellow arrowhead; **Figure S1**), suggesting that in addition to being an elongation inhibitor on some ORFs, THR acts as a termination inhibitor on the other ones. Subsequent ribosome profiling experiments can provide additional insights into the context-specificity of THR action by analyzing the transcriptome-wide distribution of ribosomes along the actively translated genes.

### Thermorubin does not interfere with the P-site tRNA

The exact location of the THR binding site in the context of the vacant 70S ribosome was known from the previous 3.2Å structure (8). While laying the foundation for understanding the mechanisms of THR action, nevertheless, this structure represents a physiologically irrelevant complex because, in the cell, antibiotics interact primarily not with vacant but with translating ribosomes associated with ligands, such as mRNAs, tRNAs, nascent protein chain and/or translation factors. Therefore, to address the possibility that the THR binding site on the ribosome is somewhat different or altered in the presence of native tRNA substrate(s), we set to determine structures of bacterial ribosome-THR complexes carrying either a single P-site tRNA or both A- and P-site tRNAs.

First, we crystallized *T. thermophilus* (*Tth*) 70S ribosomes in the presence of mRNA, initiator tRNA_i_^fMet^, and THR (**Figure 2A**), and determined its structure at 2.7Å resolution (**Figure 3A; Table S2**). The better-quality electron density maps allowed us to visualize and unambiguously model the drug (**Figure 2B, C**) as well as the 16S and 23S nucleotides adjacent to its binding site. The inclusion of mRNA and P-site tRNA in our ribosome complexes likely provided additional stabilization of the ribosome that, in turn, contributed to the ∼0.5Å higher resolution compared to the previous structure at 3.2Å (8). Most importantly, the binding position of THR in our structure is identical to the one observed previously for THR bound to the vacant *Tth* 70S ribosome in the absence of mRNA and P-site tRNA (**Figure 3B**) (8). This suggests that the presence of the initiator tRNA in the P site does not affect the general mode of THR binding to the ribosome.

**Figure 3.**
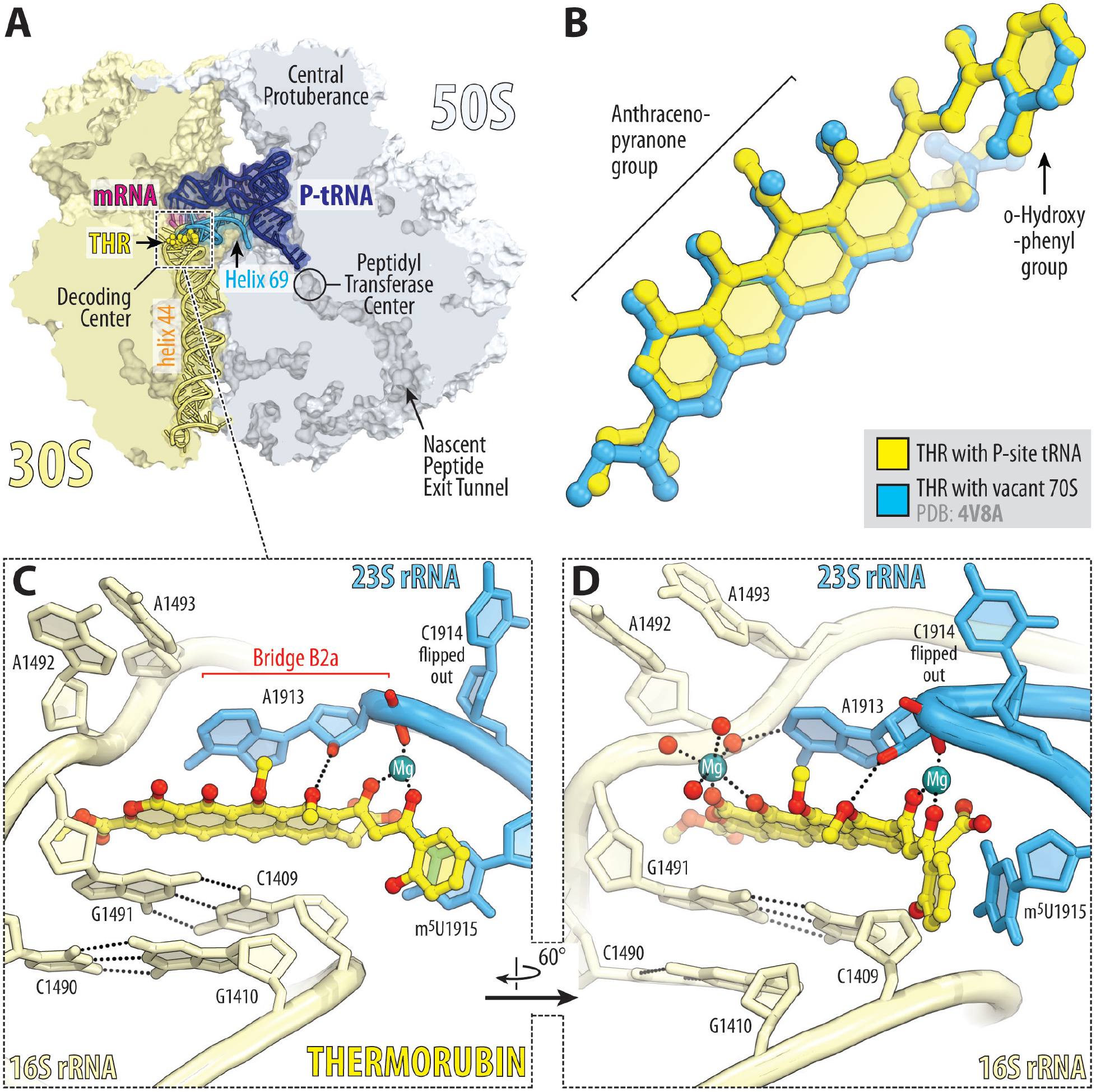
X-ray crystal structure of 70S-THR complex with P-site tRNA. (**A**) Overview of the drug-binding site (yellow) in the *T. thermophilus* 70S ribosome carrying mRNA (magenta) and deacylated initiator tRNA in the P site (navy) viewed as a cross-cut section through the nascent peptide exit tunnel. The 30S subunit is shown in light yellow; the 50S subunit is light blue. Helices 69 and 44 of the 23S and 16S rRNAs are highlighted in blue and pale yellow, respectively. (**B**) Superposition of the ribosome-bound THR in the presence of P-site tRNA (yellow) with the previous structure of THR bound to a vacant ribosome (blue, PDB entry 4V8A (8)). The structures were aligned based on helix 44 of the 16S rRNA. (**C, D**) Close-up views of the THR bound in the DC of the 70S ribosome (*E. coli* numbering of the 23S and 16S rRNA nucleotides is used). Potential H-bond interactions are indicated with dashed lines. Note that binding of THR to the 16S-23S bridge B2a causes nucleotide C1914 to flip out of its usual location in the absence of A-site tRNA. Also note that THR binding, even with the empty A site, induces nucleotides A1492 and A1493 of the 16S rRNA to flip out of helix 44.

In our structure, THR binds in the loop at the top of h44, where it forms extensive π-π stacking interactions with the surrounding nucleobases of rRNAs (**Figure 3C**). The aromatic tetracyclic moiety of the drug is sandwiched between the C1409:G1491 base pair of the 16S rRNA on one side and nucleobase A1913 of the 23S rRNA on the other (**Figure 3C**). Despite the abundance of hydrogen-bond (H-bond) donors and acceptors in the THR molecule, there is only a single H-bond that directly involves the drug with the 2’-hydroxyl of residue A1913 of the 23S rRNA. Additional electrostatic interactions of THR with the ribosome are mediated by a water-coordinated magnesium ion clearly visible in the electron density map (**Figure 2B, C; Figure 3D**). In the drug-free *Tth* 70S ribosome, nucleotide C1914, located at the tip of H69, stacks upon the next nucleotide m^5^U1915 (**Figure S2A**). However, in the presence of THR, the m^5^U1915 nucleobase stacks with the aromatic orthohydroxyphenyl moiety of the drug, thereby taking the place of C1914 and causing it to flip out (**Figure 3D; Figure S2B, C**). THR binding to the DC of the ribosome also rearranges nucleotide A1913, which becomes engaged in extensive stacking interactions with the drug and can no longer interact with the incoming A-site tRNA (**Figure S2**). Similar re-orientations of nucleotides A1913 and C1914 were observed previously in the 70S-THR structure featuring the vacant 70S ribosome (8).

### Thermorubin and A-site tRNA can co-exist on the ribosome

In principle, the flipped-out conformation of C1914 is incompatible with a tRNA in the ribosomal A site (**Figure S2B**), as suggested before (8). However, our toe-printing data contradict this hypothesis and indicate that the ribosome is able to translate ORFs in the presence of THR (**Figure 1C, D; Figure S1**) – a result that could hardly be reconciled if THR is incompatible with an accommodated A-site tRNA. This observation suggests that, in the presence of THR, accommodation of aa-tRNA into the ribosomal A site either causes re-orientation of nucleotide C1914 and pushes it out of the way, or repositions the drug into a previously unseen conformation, or both.

Therefore, in order to directly address the possibility of A-tRNA-induced C1914 re-orientation, we determined the 2.7Å-resolution cryo-electron microscopy (cryo-EM) structure of the *E. coli* 70S ribosome carrying Phe-tRNA^Phe^ and tRNA_i_^fMet^ in the A and P sites, respectively, and also bound with THR (**Figure 4A; Table S3**). Contrary to the previous hypothesis and *in silico* predictions (**Figure S2B**), our cryo-EM structure reveals fully accommodated Phe-tRNA^Phe^ and THR both bound to the ribosome side-by-side (**Figure 4A, B; Figure S3**). Interestingly, in the presence of THR, we observed a ∼90-degree deflection of nucleotide C1914 towards the solvent side relative to its flipped-out conformation seen in the previous structure, allowing Phe-tRNA^Phe^ to fully accommodate into the A site (**Figure 4B, C**). At the same time, most of the interactions of THR with the ribosome were retained in the presence of the A-site tRNA, and its binding site remained undisturbed (**Figure 4C**). This structure clearly shows that an aa-tRNA can accommodate into the ribosomal A site even in the presence of ribosome-bound THR, rationalizing our *in vitro* toe-printing data (**Figure 1C, D**).

**Figure 4.**
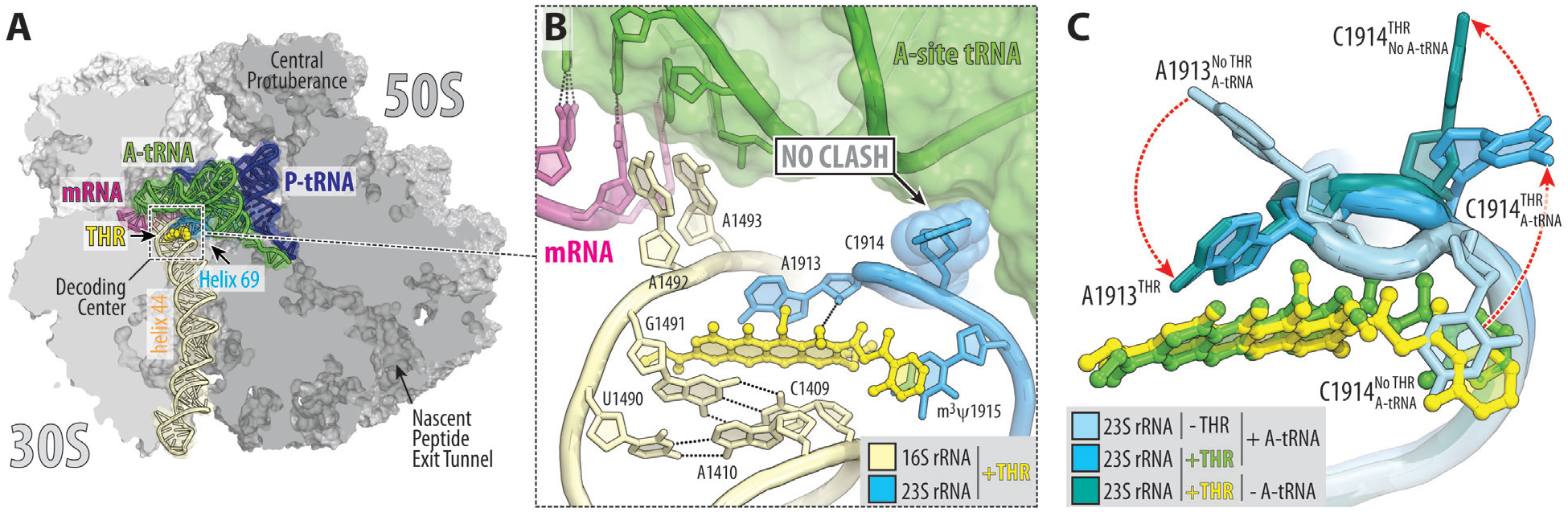
Cryo-EM structure of 70S-THR complex with both A- and P-site tRNAs. (**A**) Overview of the THR binding site (yellow) in the *E. coli* 70S ribosome containing mRNA (magenta) and full-length Phe-tRNA^Phe^ (green) and tRNA_i_^fMet^ (navy) in the A and P sites, respectively, viewed as a cross-cut section through the ribosome. (**B**) Close-up view of THR bound in the DC in the presence of fully accommodated A-site tRNA. Note that nucleotide C1914 adopts a previously unseen conformation, in which it does not interfere with tRNA binding. (**C**) Structural rearrangements in the DC (red dashed arrows) upon THR binding and tRNA accommodation. Superpositioning of the previous structure of drug-free ribosome carrying both A- and P-site tRNAs (PDB entry 6XHW (23)) with the two new 70S-THR structures with or without A-site tRNA. The structures were aligned based on helix 44 of the 16S rRNA. Note that in the presence of THR and A-site tRNA, nucleotide C1914 flips halfway out from its normal position in Helix 69 of the 23S rRNA, allowing for tRNA accommodation (light blue to blue transition). However, in the absence of A-site tRNA, nucleotide C1914 is unrestricted and fully flips out (blue to teal transition).

### Thermorubin affects multiple steps of translation elongation

Altogether our toe-printing and structural data suggest that THR does not inhibit translation initiation and binding of tRNA to the ribosomal A site. However, the observed ribosome stalling within the ORF of mRNA templates used for toe-printing (**Figure 1C, D**, lane 2, blue, green, red, teal, and orange arrowheads) suggests that one or more steps of elongation could be impaired by THR. As the elongation cycle consists of (i) A-site tRNA delivery, (ii) peptide bond formation, and (iii) translocation steps, we set to assess the effects of THR on each of these individual steps using a reconstituted *in vitro* translation system. First, to monitor the effect of THR on tRNA delivery, we used a stopped-flow fluorescence detection assay to follow the delivery of Phe-tRNA^Phe^(Prf16/17) in the form of the ternary complex (EF-Tu•GTP•Phe-tRNA^Phe^(Prf16/17)) to the initiated ribosome complexes bearing the UUU codon in the vacant A site (**Figure 5A**). An increase in a characteristic biphasic fluorescence change represents the initial binding of the ternary complex to the ribosome and subsequent codon recognition, which induces GTPase activation and results in GTP hydrolysis by EF-Tu. The subsequent decay of the high fluorescence intermediate is related to the release of tRNA from the GDP-bound form of EF-Tu and accommodation of the tRNA in the A site of the ribosome (49). Two-exponential fitting of the fluorescence curves allowed us to measure the rate constants of initial ternary complex binding and subsequent tRNA accommodation. Interestingly, THR does not affect ternary complex binding (k_app1_(No Drug) = 22.8 ± 0.9 s^−1^ *vs*. k_app1_(THR) = 19.0 ± 0.4 s^−1^), whereas the rate of tRNA accommodation decreases by approximately eight folds (k_app2_(No Drug) = 9.4 ± 0.3 s^−1^ *vs*. k_app2_(THR) = 1.19 ± 0.04 s^−1^) suggesting that THR slows down aa-tRNA accommodation.

**Figure 5.**
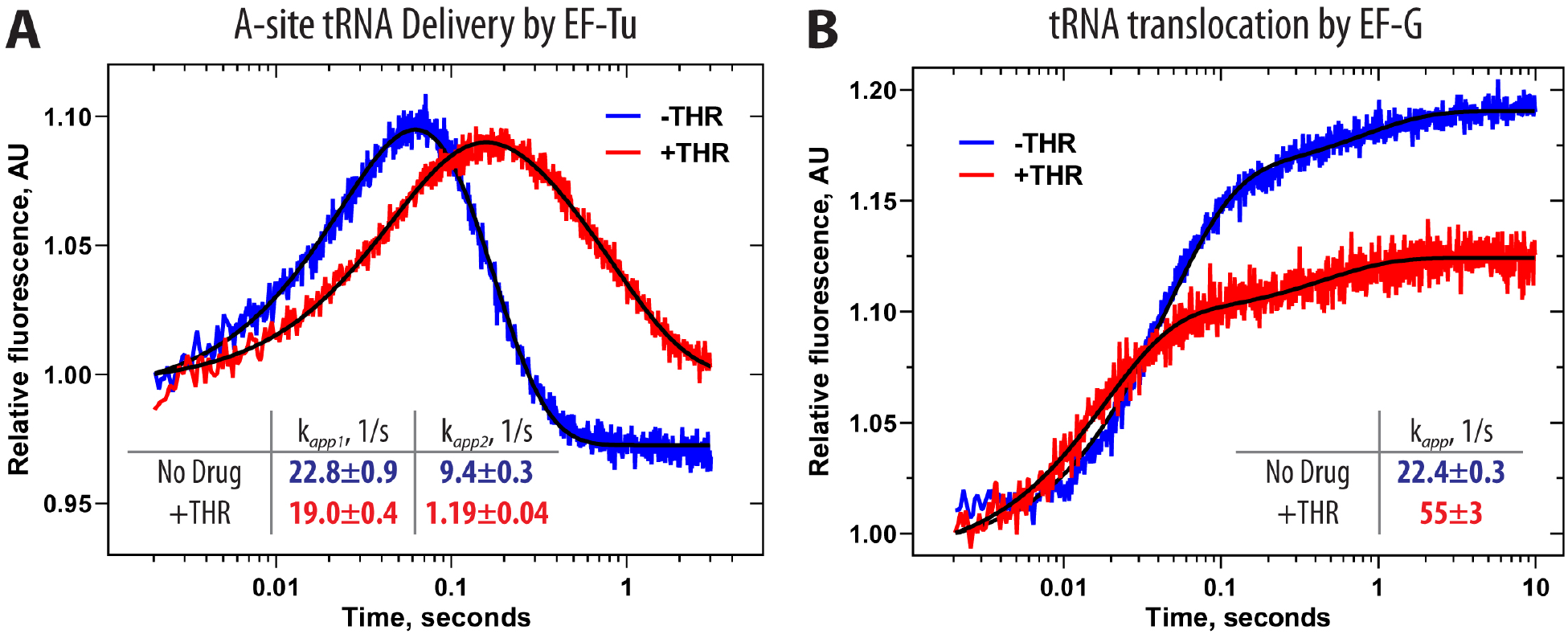
Ribosome-bound THR impairs the elongation cycle. (**A**) Pre-steady state kinetics of A-site tRNA binding upon interaction of the ternary complex EF-Tu•GTP•Phe-tRNA^Phe^(Prf16/17) (0.1 µM) with the 70S initiated ribosomes (0.4 µM) containing fMet-tRNA_i_^fMet^ in the P site. Shown are time courses in the absence (blue) and presence (red) of THR. (**B**) Pre-steady state kinetics of translocation upon interaction of the 70S pre-translocation ribosome complexes (60 nM) containing deacylated-tRNA_i_^fMet^ in the P site and fMet-Phe-tRNA^Phe^(Prf16/17) in the A site with EF-G (2 µM). Color scheme is the same as in panel A. Each time course represents the average of 5 to 7 experimental replicates. Standard deviations associated with the kinetics were calculated using GraphPad Prism software.

Accommodation of the aa-tRNA into the A site of the PTC leads to peptide bond formation by spontaneous transfer of a peptidyl moiety of the P-site tRNA onto the A-site tRNA. Thus, the final level of dipeptide formed depends on the amount of correctly accommodated A-site tRNA. We found that THR-containing ribosomal complexes are less effective in the formation of fMet-[^14^C]Phe-dipeptidyl-tRNA^Phe^, with only 59% of dipeptide formed relative to the drug-free ribosomes implying that THR affects tRNA accommodation.

Peptide bond formation is followed by a translocation event that comprises large-scale movements of mRNA and tRNAs through the ribosome, marking the end of each elongation cycle. During this step, the deacylated-tRNA moves from the P to the E site, which then rapidly dissociates from the ribosome. Simultaneously, peptidyl-tRNA translocates from the A to the P site maintaining codon-anticodon interactions with the mRNA that results in ribosome movement by one codon along mRNA. To assess whether THR affects translocation, we monitored the kinetics of this process by changes in fluorescence upon addition of the translational GTPase EF-G to the pre-translocation ribosome complex containing fluorescently labeled fMet-Phe-tRNA^Phe^(Prf16/17) in the A site and deacylated tRNA_i_^fMet^ in the P site. The addition of EF-G to such complex results in a two-step change in fluorescence intensity with a predominant fast step (more than 80% of the total signal amplitude). The addition of THR increases the rate of peptidyl-tRNA translocation from the A to the P site by 2.5 fold (k_No Drug_ = 22.4 ± 0.3 s^-1^ *vs*. k_THR_ = 55 ± 3 s^-1^), whereas the amplitude of the total fluorescence signal decreases by 30% (**Figure 5B**). Both of these observations indicate that A-site tRNA becomes destabilized in the presence of THR (similar to (46,50)), resulting either in dissociation of the dipeptidyl-tRNA from the ribosome before translocation or in its more rapid translocation to the P site upon addition of EF-G. These data agree with the observed slowed-down accommodation of aa-tRNA, suggesting that, although THR does not preclude tRNA binding to the ribosomal A site, THR-induced re-arrangements of rRNA nucleotides are likely responsible for the observed poor stability of the A-site tRNA. Indeed, our cryo-EM data reveals that in the presence of THR, nucleotide A1913 of the 23S rRNA is unable to form H-bond with the 2’-OH of the ribose of nucleotide 37 in the A-site tRNA rationalizing the lower stability of A-site tRNA.

### Thermorubin decreases the binding affinity of tRNA to the A site

Purine nucleotide at position 37 in most tRNAs is usually heavily post-transcriptionally modified and plays a crucial role in maintaining the structure of the tRNA anticodon loop and stability of the codon-anticodon duplex (46). Removal of the modification or replacement of purine at this position with pyrimidine dramatically decreases the enthalpy of tRNA-ribosome interactions (46), suggesting that the observed loss of H-bond between this nucleotide and 23S rRNA due to THR-induced rearrangement of A1913 (**Figure S2A, B**) might influence the thermodynamic properties of A-site tRNA binding. To evaluate the effect of THR on the affinity of peptidyl-tRNA to the A site, we determined dissociation constants (46). Under appropriate conditions, A-site peptidyl-tRNA can dissociate reversibly from the purified pre-translocation ribosome complexes depending on its binding stability (51), allowing equilibrium dissociation constant (K_d_) and the rate constants of tRNA dissociation from (k_off_) and association with the A site (k_on_) to be calculated. When the drug-free ribosome complexes containing A-site peptidyl-tRNA^Val^ were incubated at 37°C and 10 mM Mg^2+^, approximately 50% of the peptidyl-tRNA remained bound to the ribosomal A site even after 2 hours of incubation (**Figure 1E**, blue trace). In contrast, dissociation of the same amount of peptidyl-tRNA^Val^ from the THR-containing ribosome complexes occurred significantly faster, leaving only about 10% of the peptidyl-tRNA bound to the ribosomes after 10 minutes of incubation (**Figure 1E**, red trace). The calculated affinity of peptidyl-tRNA^Val^ to the A site of the THR-bound ribosome is 12 times lower compared to the drug-free ribosome complexes (K_d_(No drug) = 0.13 ± 0.08 µM *vs*. K_d_(THR) = 1.5 ± 0.3 µM) (**Figure 1E; Table S4**), yet again emphasizing that A-site tRNA binding is negatively affected by THR.

### Ribosome-bound thermorubin likely interferes with the activity of class-I RFs

THR-dependent ribosome stalling at the stop codons of *rst1, rst2, ermBL*, and *yrbA* mRNA templates used in our toe-printing assay (**Figure 1C, D; Figure S1**) suggests that THR can also act as an inhibitor of translation termination. In the light of this finding, we explored whether THR simply prevents binding of class-I release factors (RF1 and/or RF2) to the A site or, *vice versa*, traps class-I RFs on the ribosome, similar to the mode of action of the peptide antibiotic apidaecin (52), thereby making translation termination impossible in either case. To distinguish between these two possibilities, we attempted to obtain structures of the 70S-RF1 complex with THR using ribosomes either from *T. thermophilus* or *E. coli* and using either X-ray crystallography or cryo-EM approaches. Despite numerous attempts, our structures showed that RF1 did not bind to the 70S-THR complexes, suggesting that THR sterically hinders the delivery of RF1 into the A site. Superimposition of the 70S-THR structure with the previous structures of ribosome-bound RF1 (**Figure 6A**) reveals a relatively small sterical hindrance between the ribosome-bound drug and Arg286 residue of the RF1 bound in the A site. Although even a small sterical overlap between the drug and RF1 should prevent binding of either one to the ribosome, the Arg286 residue does not interact with anything, is not restricted by the surrounding residues and, in principle, shall be able to reorient to avoid the clash with THR. However, the THR-induced flipped-out conformation of nucleotide C1914 of the 23S rRNA produces extensive sterical clash with residues Glu110, Arg112, Lys157, and Ile293 of the A-site-bound RF1, suggesting that RF is unlikely to be able to accommodate into the A site in the presence of THR (**Figure 6A**) explaining why we were unable to obtain structures of such complexes. Moreover, binding of RF to the ribosome induces significant rearrangements of nucleotides A1492, A1493 of the 16S rRNA and A1913 of the 23S rRNA, which, in turn, are incompatible with THR (**Figure 6B**). Altogether our structural analysis shows that although binding positions of THR and class-I RFs on the ribosome barely overlap, binding of one of these ligands to the ribosome induces mutually exclusive conformations of nucleotides around the DC, preventing binding of the other.

**Figure 6.**
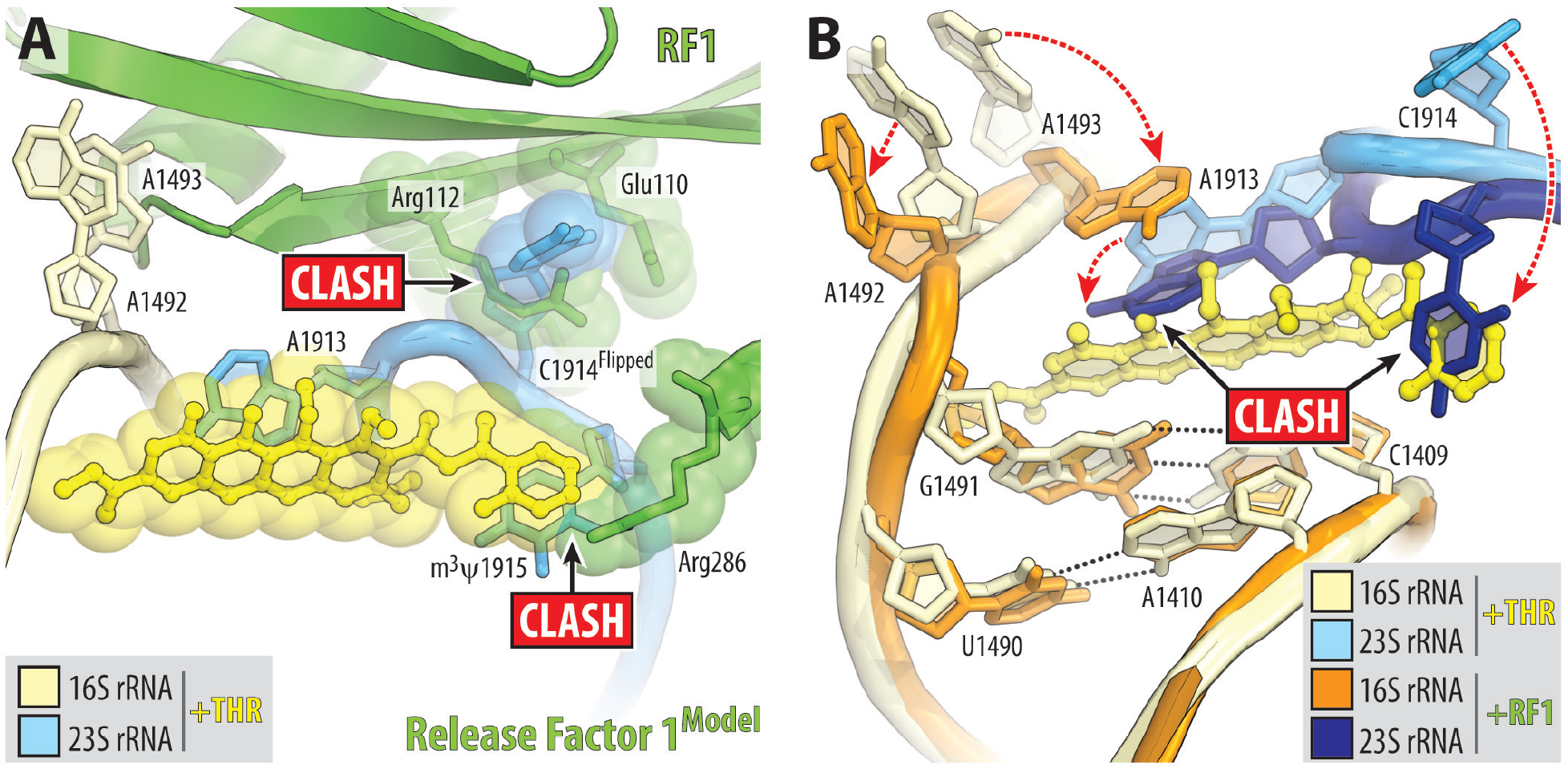
Ribosome-bound THR likely prevents binding of release factor to the A site. (**A**) Superimposition of the ribosome-bound RF1 (PDB entry 3D5A (54)) with the new structure of 70S-THR complex carrying both A- and P-site tRNAs. The structures were aligned based on helix 44 of the 16S rRNA. Note that the THR-induced flipped-out conformation of C1914 would cause steric clashes with RF1 bound in the A site. (**B**) Structural re-arrangements in the DC that are required for RF1 to bind to the 70S ribosome (dashed red arrows). Note that binding of RF to the ribosome requires nucleotides A1492, A1493 of the 16S rRNA and A1913 of the 23S rRNA to adopt positions that are incompatible with ribosome-bound THR.

## CONCLUSION

Altogether, our data suggest that THR is a potent inhibitor of protein synthesis that targets the elongation and/or termination steps of translation. Interestingly, the effects of THR binding on translation resemble those observed from mutations in bridge B2a (53), yet again suggesting interference with the functioning of bridge B2a to be the main mode of action of THR. Despite its strong antimicrobial properties, THR is not currently used in the clinic mainly due to its poor solubility and low thermostability above 37°C (5). Chemical derivatizations of the drug could help improve its physicochemical (solubility) and pharmacokinetic (bioavailability) properties. It is intriguing, however, that a compound with such poor solubility and low thermostability is naturally produced by a bacterial host whose optimal growth temperature is between 48-53°C (5). This makes us wonder if THR is complexed with an adjuvant, which stabilizes and solubilizes it, in the cytoplasm of the producing bacterial cells. Perhaps, an alternate purifying strategy might help the isolation of THR in its native form, which may be more soluble and/or potent than the parent compound.

## DATA AVAILABILITY STATEMENT

Coordinates and structure factors were deposited in the RCSB Protein Data Bank under the accession code **7XXX** for the *T. thermophilus* 70S ribosome crystal structure in complex with THR, mRNA, and deacylated P-site **tRNA**_**ifMet**_. The cryo-EM map for the *E. coli* 70S ribosome in complex with THR, mRNA, aminoacylated A-site **Phe-tRNA**^**Phe**^, and deacylated P-site **tRNA**_**ifMet**_ was deposited in the Electron Microscopy Data Bank (EMDB) under the accession code **EMD-YYYY**, with the coordinates deposited in the RCSB Protein Data Bank under the accession code **7ZZZ**.

All previously published structures that were used in this work for structural comparisons were retrieved from the RCSB Protein Data Bank: PDB entries 4V8A, 6XHW, and 3D5A.

No sequence data were generated in this study.

## FUNDING

This work was supported by the National Institutes of Health [R01-GM132302 to Y.S.P.; R01-GM136936 to M.G.G.], the Welch Foundation [H-2032-20200401 to M.G.G.], the Illinois State startup funds [to Y.S.P.], Russian Ministry of Science [075-15-2021-1396 to V.I.M.], and Russian Science Foundation [22-14-00278 to A.L.K., pre-steady state kinetics and thermodynamics of elongation reactions]. Funding for open access charge: National Institutes of Health [R01-GM132302 to Y.S.P.; R01-GM136936 to M.G.G.] and Russian Science Foundation [22-14-00278 to A.L.K.].

## CONFLICT OF INTEREST STATEMENT

The authors declare no competing financial or non-financial interests.

## AUTHOR CONTRIBUTIONS

MNP performed *in vitro* inhibiton assay; VIM and TPM performed dual-reported assay; MNP, VIM, and TPM performed toe-printing assays on various mRNA templates; MNP and YSP performed X-ray crystallography studies; MGG performed cryo-EM data analysis and structure determination; AAG, OAT, and EP performed pre-steady state kinetics and thermodynamics studies; YSP, MGG, ALK, PVS, and IAO supervised the experiments. All authors interpreted the results. MNP, AP, ALK, MGG, and YSP wrote the manuscript.

## ACKNOWLEDGMENTS

We thank Dr. Mariia Rybak for preparing the 70S-THR ribosome complex bound to Phe-tRNA^Phe^ in the A site and tRNA_i_^fMet^ in the P site, and for collecting and processing cryo-EM data. We thank Dr. Francis Johnson for providing thermorubin. We thank the staff at NE-CAT beamlines 24ID-C and 24ID-E for help with X-ray diffraction data collection, especially Drs. Malcolm Capel, Frank Murphy, Igor Kourinov, Anthony Lynch, Surajit Banerjee, David Neau, Jonathan Schuermann, Narayanasami Sukumar, James Withrow, Kay Perry, Ali Kaya, and Cyndi Salbego. We are thankful to Dr. Michael Sherman for help with cryo-EM data acquisition, the Sealy Center for Structural Biology and Molecular Biophysics of the University of Texas Medical Branch at Galveston for providing critical infrastructure and expertise, and Drs. Ka-Yiu Wong and John Perkyns for computational support.

This work is based upon research conducted at the Northeastern Collaborative Access Team beamlines, which are funded by the National Institute of General Medical Sciences from the National Institutes of Health [P30-GM124165 to NE-CAT]. The Eiger 16M detector on 24ID-E beamline is funded by an NIH-ORIP HEI grant [S10-OD021527 to NE-CAT]. This research used resources of the Advanced Photon Source, a U.S. Department of Energy (DOE) Office of Science User Facility operated for the DOE Office of Science by Argonne National Laboratory under Contract No. DE-AC02-06CH11357. This work used UCSF Chimera for visualization of cryo-EM volume maps and initial structure modeling [P41-GM103311 to the Resource for Biocomputing, Visualization, and Informatics at the University of California, San Francisco].

## Notes

### Competing Interest Statement

The authors have declared no competing interest.

