## Supplementary Information for "Insights into the molecular mechanism of translation inhibition by the ribosome-targeting antibiotic thermorubin"

**This file includes:**

I. Supplementary Tables 1 to 4;
II. Supplementary Figures 1 to 5 with legends;
III. Supplementary References.

### II. SUPPLEMENTARY TABLES

#### Table S1. DNA primers used for generating the templates for toe-printing analysis.

| Toe-printing template | Primer name | Used for | Nucleotide sequence (5'-to-3') |
|---|---|---|---|
| *rst1* | RST1-Fwd | PCR | TAATACGACTCACTATAGGGCTTAAGTATAAGGAGGAAAACATATGAAATTCGCCATCACCCTGCGTCAGTGCGAAGGCTGGTCACC |
| | RST1-Rev | PCR | GGTTATAATGAATTTTGCTTATTAACGATAGAATTCTATCACTTTTTTTATTATTAATTGTCATAATGTACCGGTGACCAGCCTTCGCAC |
| *rst2* | RST2-Fwd | PCR | |
| | RST2-Rev | PCR | |
| *ermBL* | ErmB3-Fwd | PCR | TAATACGACTCACTATAGGGCTTAAGTATAAGGAGGAAAAAATATGTTGGTATTCCAAATGCGTAATGTAGATAAAACATCTAC |
| | ErmB3-Rev | PCR | GGTTATAATGAATTTTGCTTATTAACGATAGAATTCTATCACTTATTTCAAAATAGTAGATGTTTTATCTACATTACG |
| *yrbA* | YRBA-Fwd | PCR | GGTTATAATGAATTTTGCTTATTAACTCGACATCGCATCAGGATTCAGCACGTGAATCCAACTGTCGAGGTCG |
| | YRBA-Rev | PCR | GTATAAGGAGGAAAACATATGATATACCCCTGCGGAGTGGGCGCGCGATCGCAAACTGAACGGCTTTAGGCCGACCTCGACAGTTGGAT |
| | NV1 | Toe-printing | GGTTATAATGAATTTTGCTTATTAAC |

**Table S2. X-ray data collection and refinement statistics.**

| ***Crystals*** | | ***T. thermophilus* 70S ribosome in complex with mRNA, $tRNA_i^{Met}$, and THR PDB entry 7XXX** |
|---|---|---|
| ***Diffraction data*** | | |
| Space Group | | $P2_12_12_1$ |
| Unit Cell Dimensions, Å (a x b x c) | | 209.51 x 449.73 x 622.15 |
| Wavelength, Å | | 0.9792 |
| Resolution range (outer shell), Å | | 190-2.70 (2.77-2.70) |
| I/σI (outer shell) | | 8.87 (1.00) |
| Resolution at which I/σI=1, Å | | 2.70 |
| Resolution at which I/σI=2, Å | | 2.90 |
| CC(1/2) at which I/σI=1, % | | 17.7 |
| CC(1/2) at which I/σI=2, % | | 50.0 |
| Completeness (outer shell), % | | 97.0 (95.9) |
| $R_{merge}$ (outer shell)% | | 16.6 (177.4) |
| No. of crystals used | | 1 |
| No. of Reflections Used: | Observed | 8,236,656 |
| | Unique | 1,538,795 |
| Redundancy (outer shell) | | 5.35 (5.18) |
| ***Refinement*** | | |
| Resolution range of the diffraction data included in the refinement, Å | | 155-2.70 |
| $R_{work}/R_{free}$, % | | 20.3/25.5 |
| *No. of Non-Hydrogen Atoms* | | |
| RNA | | 194,201 |
| Protein | | 90,876 |
| Ions (Mg, K, Zn, Fe) | | 2,795 |
| Waters | | 4,325 |
| *Ramachandran Plot* | | |
| Favored regions, % | | 88.50 |
| Allowed regions, % | | 9.37 |
| Outliers, % | | 2.12 |
| *Deviations from ideal values (RMSD)* | | |
| Bond, Å | | 0.009 |
| Angle, degrees | | 1.428 |
| Chirality | | 0.059 |
| Planarity | | 0.007 |
| Dihedral, degrees | | 17.296 |
| Average B-factor (overall), $Å^2$ | | 57.5 |

**Table S3. Cryo-EM data collection, refinement, and validation statistics.**

| ***Dataset*** | ***E. coli* 70S ribosome in complex with mRNA, A-site Phe-tRNA$^{Phe}$, P-site tRNA$_i^{Met}$ and THR PDB entry 7XXX EMDB entry XXXX** |
|---|---|
| ***Data collection and processing*** | |
| Magnification | 96,000x |
| Voltage (kV) | 300 |
| Electron exposure (e$^-$/Å$^2$) | 40 |
| Defocus range (μm) | –1 to –2.3 |
| Pixel size (Å) | 0.86 |
| Symmetry imposed | C1 |
| Initial particle images (no.) | 982,190 |
| Final particle images (no.) | 310,702 |
| Map resolution (Å) | 2.7 |
| FSC threshold | 0.143 |
| ***Refinement*** | |
| Initial model used (PDB code) | 6GXP |
| Model resolution (Å) | 2.9 |
| FSC threshold | 0.5 |
| Map sharpening B factor (Å$^2$) | –80 |
| $CC_{mask}$ | 0.88 |
| MolProbity score | 1.91 |
| Clashcore | 6.25 |
| *No. of Non-Hydrogen Atoms* | |
| RNA | 99,952 |
| Protein | 45,349 |
| Ions (Mg, Zn) | 561 |
| Waters | 299 |
| *Ramachandran Plot* | |
| Favored regions, % | 89.21 |
| Allowed regions, % | 10.39 |
| Outliers, % | 0.40 |
| *Deviations from ideal values (RMSD)* | |
| Bond, Å | 0.007 |
| Angle, degrees | 0.829 |
| Chirality | 0.049 |
| Planarity | 0.007 |
| Dihedral, degrees | 14.755 |
| Average B-factor (overall), Å$^2$ | 57.9 |

**Table S4. Thermodynamic and kinetic parameters of peptidyl-tRNA binding in the A site.**

| ***70S Ribosome Complex*** | **No Drug** | **+THR** |
|---|---|---|
| $K_d$, μM | 0.13 ± 0.08 | 1.5 ± 0.3 |
| $k_{off}$, $10^{-3}\cdot sec^{-1}$ | 0.46 ± 0.08 | 3.6 ± 0.7 |
| $k_{on}$, $10^{3}\cdot M^{-1}sec^{-1}$ | 3.5 ± 0.6 | 2.4 ± 0.5 |

### II. SUPPLEMENTARY FIGURES

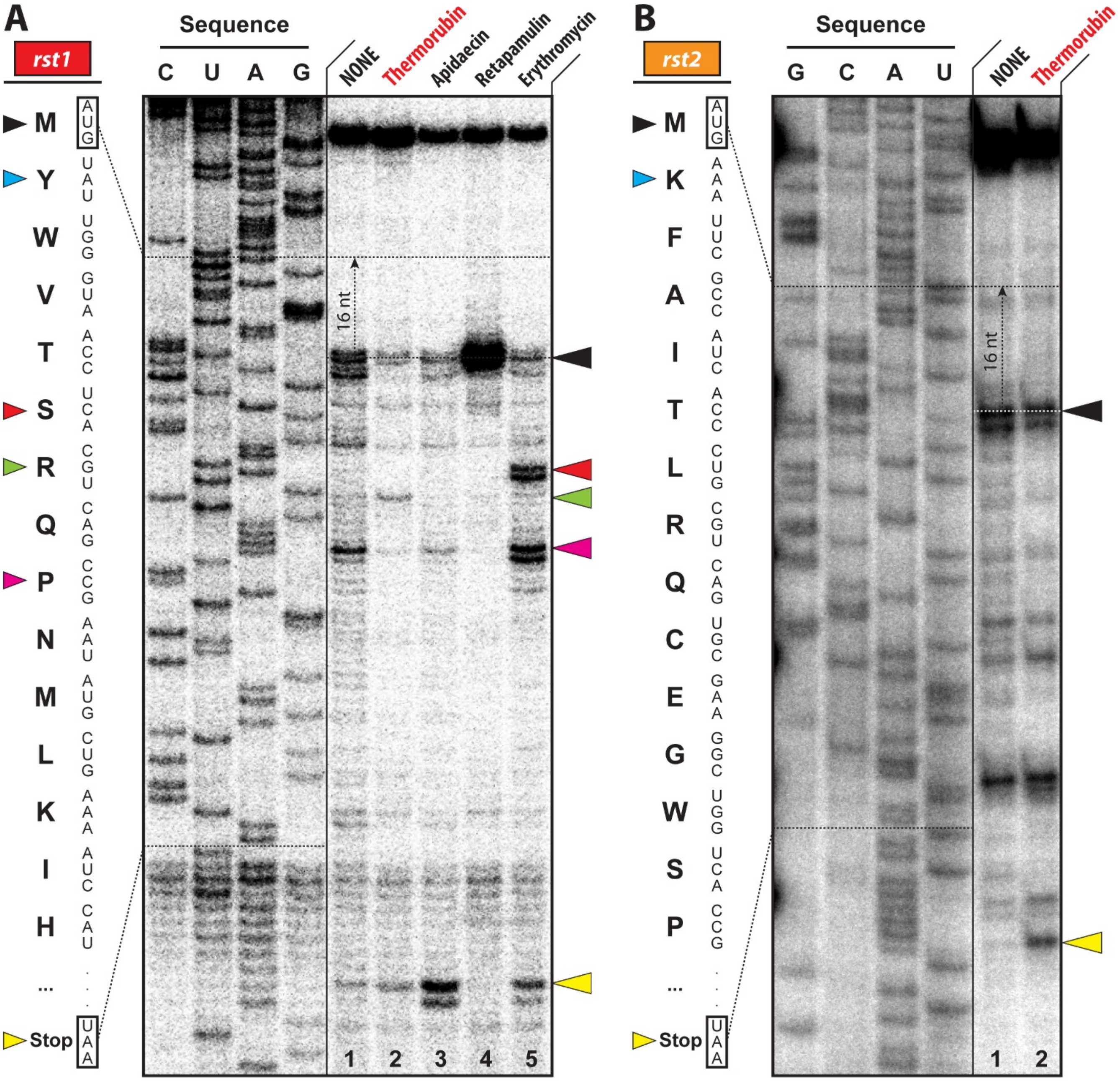


**Figure S1. Toe-printing in the presence of THR on *rst1* and *rst2* mRNA templates.** Ribosome stalling by THR on *rst1* (A) and *rst2* (B) mRNAs revealed by reverse transcription inhibition (toe-printing) in a recombinant cell-free translation system. For details, see the legend for **Figure 1C, D**. Black arrowhead marks translation start site (retapamulin control, lane 4). The green arrowhead points to the THR-induced arrest sites within the coding sequence of *rst1* mRNA. Red and magenta arrowheads point to the erythromycin-specific stalling sites in *rst1* only. Yellow arrowheads point to the translation stop site (apidaecin control, lane 3).

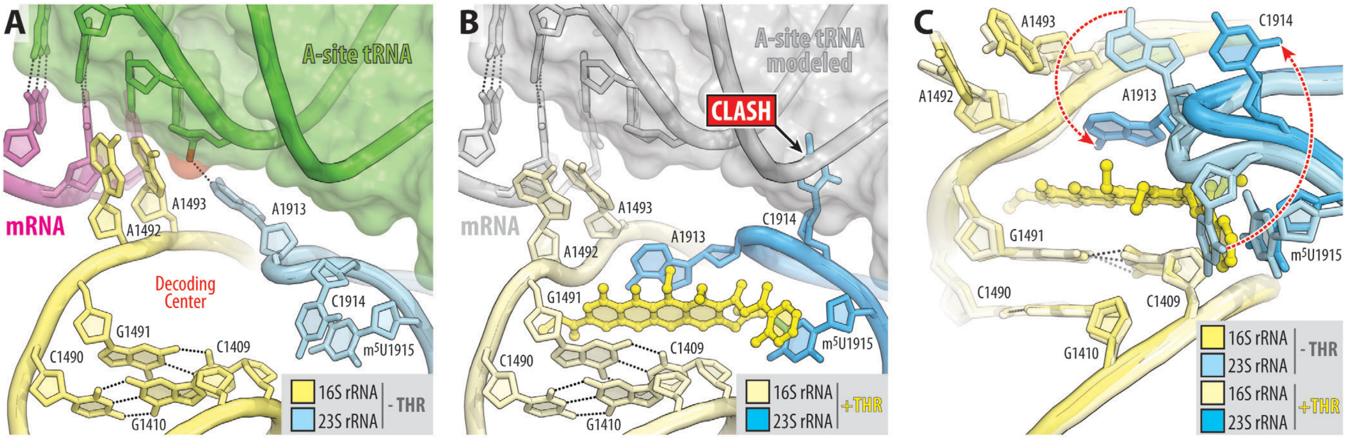


**Figure S2. Structural rearrangements in the decoding center upon THR binding to bridge B2a.** (**A, B**) Close-up views of the decoding center of the drug-free (A, PDB entry 6XHW (1)) *versus* THR-bound (B) *Tth* 70S ribosome. 16S and 23S rRNAs are colored in shades of light yellow and light blue, respectively, as indicated by the insets. In panel (A), mRNA and A-site tRNA are shown in magenta and green, respectively. In panel (B), both mRNA and tRNA are shown in grey to emphasize that THR and A-site tRNA might not be able to co-exist due to a clash between the residue C1914 of the 23S rRNA and A-site tRNA. (**C**) Superimposition of the 70S-THR structure with that of the drug-free ribosome carrying both A- and P-site tRNAs (PDB entry 6XHW (1)). The structures were aligned based on helix 44 of the 16S rRNA. Note that in the presence of THR, nucleotide A1913 relocates towards the drug and engages in extensive stacking interactions with it, whereas nucleotide C1914 flips out from its normal position in Helix 69 of the 23S rRNA (red dashed arrows).

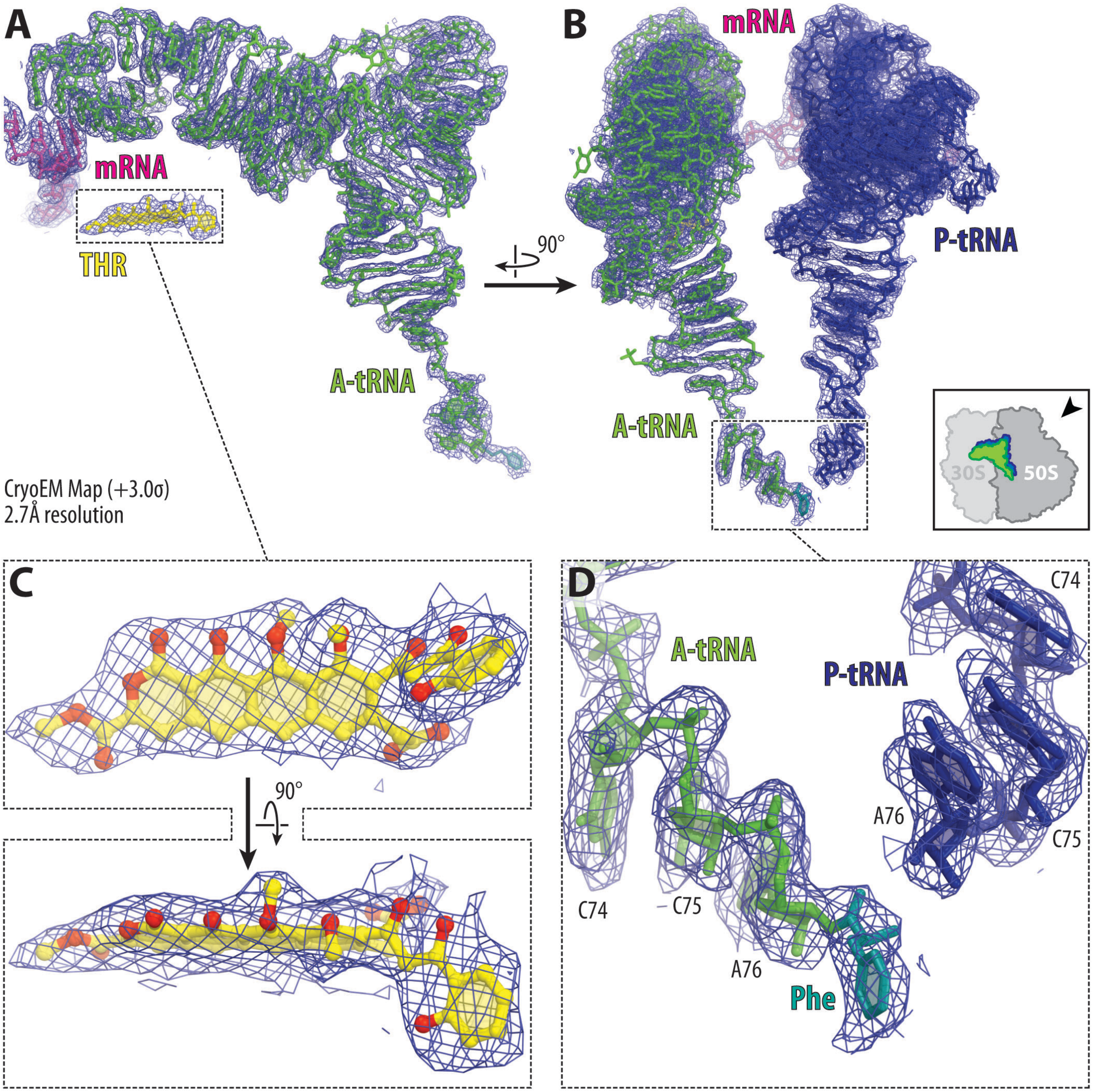


**Figure S3. Representative cryo-EM map for the 70S-THR complex with bound A- and P-site tRNAs.** (**A, B**) Cryo-EM map (blue mesh) of the ribosome-bound THR (yellow), mRNA (magenta), A- and P-site tRNAs (green and navy, respectively). The refined models of tRNAs are displayed within their respective charge density maps contoured at 3.0σ. In (B), the entire bodies of the A- and P-site tRNAs are viewed from the back of the 50S subunit, as indicated by the inset. Ribosome subunits are omitted for clarity. (**C, D**) Close-up views of the THR binding site in the DC of the 30S ribosomal subunit (C) and CCA-ends of the A- and P-site tRNAs in the PTC on the 50S ribosomal subunit. Note that, while the P-site tRNA is deacylated, the A-site tRNA is aminoacylated with phenylalanine(teal).

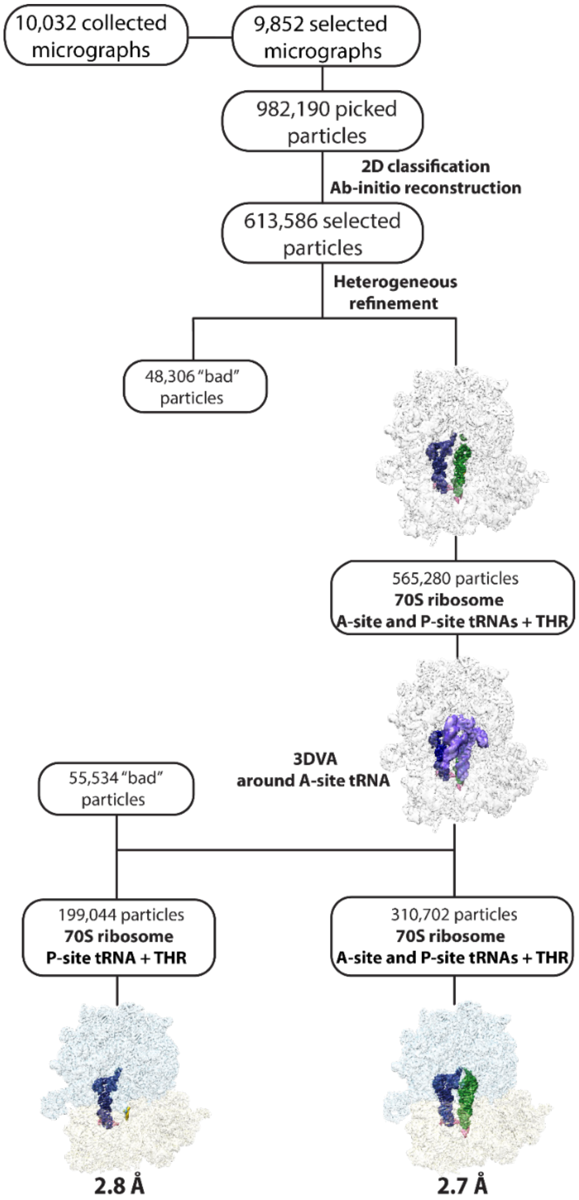


**Figure S4. Cryo-EM data processing and particle classification workflow.** All data processing steps were performed in cryoSPARC 3.3.2 (2). 10,032 micrographs were collected, of which 9,852 were selected for further processing. Two rounds of reference-free 2D classification were performed, and the selected particles were used to generate the *ab-initio* volume. 'Heterogeneous refinement' was performed to sort particles from the *ab-initio* 3D volume into five groups, allowing to discard 48,306 particles and combine two groups into one class average containing A- and P-site tRNAs (565,280 particles). Using 3D variability analysis (3DVA) focused around the A-site tRNA (semi-transparent mask shown in violet), 310,702 particles were separated for containing solid density for both A- and P-site tRNAs, while 199,044 particles contained only P-site tRNA. This process allowed to discard 55,534 particles with weak density for tRNAs. Non-uniform and CTF refinement of each class yielded reconstructions of the *E. coli* 70S ribosome with A-site Phe-$tRNA^{Phe}$, P-site $tRNA_i^{fMet}$, and THR, and with only P-site $tRNA_i^{fMet}$ and THR at nominal resolutions of 2.7Å and 2.8Å, respectively.

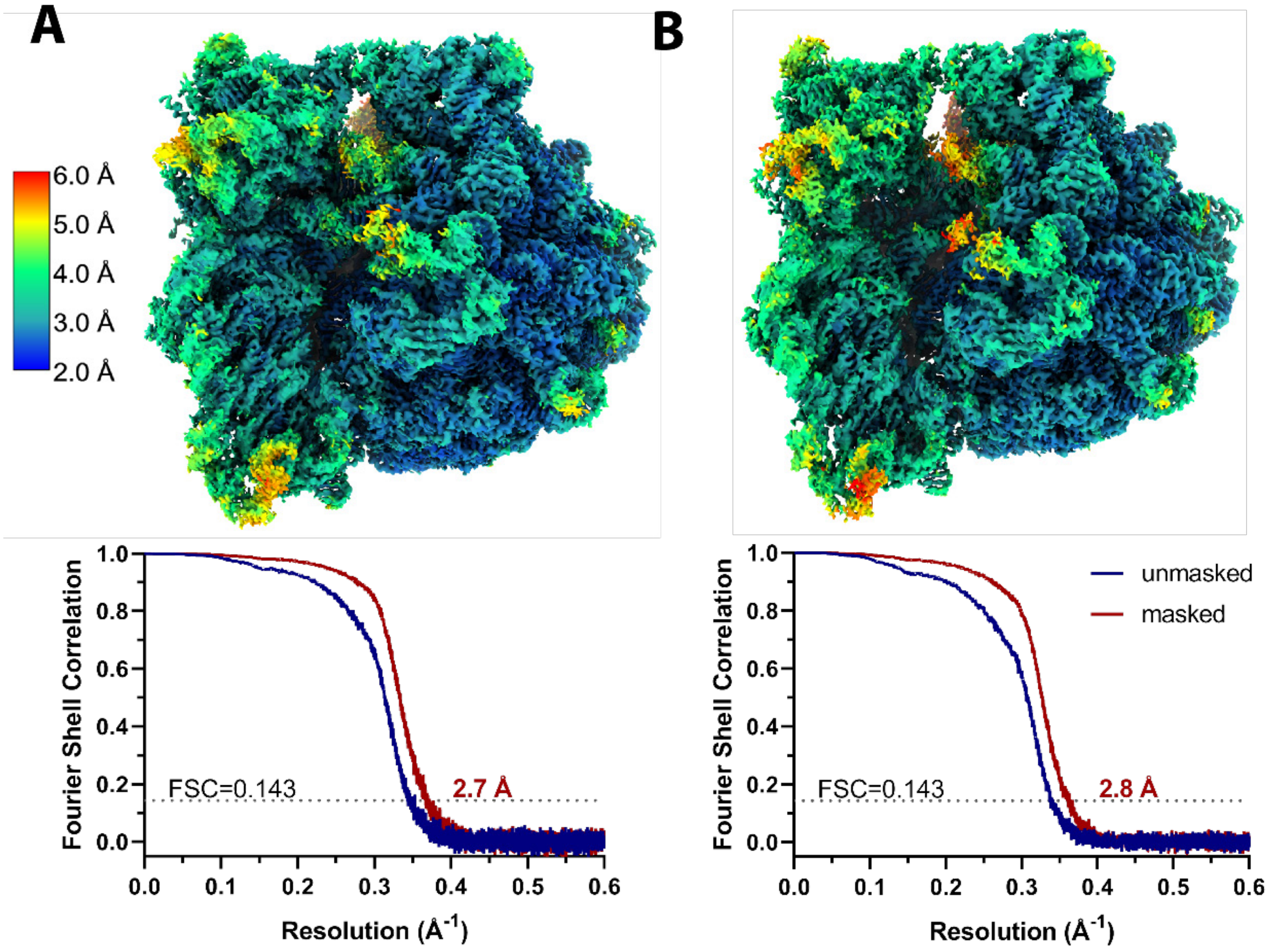


**Figure S5. Local resolution estimation and Fourier Shell Correlation (FSC) validation.** (**A, B**) Local resolution heat maps of the *E. coli* 70S ribosome with (**A**) A-site Phe-tRNA$^{Phe}$ and P-site tRNA$_i^{fMet}$ + THR, and (**B**) P-site tRNA$_i^{fMet}$ + THR shown in the range of 2 – 6Å resolution, calculated with cryoSPARC 3.3.2 implementation of BlocRes (3). Below each reconstruction, gold-standard Fourier Shell Correlation (FSC) curves of each half-map using a 'soft mask' excluding solvent (red) are plotted across resolution. Validation of the maps was performed in PHENIX 1.19.2 (4).
